# Deep Learning Assisted Smartphone-based Quantitative Microscopy for Label-free Hematological Analysis

**DOI:** 10.1101/2023.01.24.525176

**Authors:** Bingxin Huang, Lei Kang, Victor T. C. Tsang, Claudia T. K. Lo, Terence T. W. Wong

## Abstract

Hematologists evaluate alterations in blood cell enumeration and morphology to confirm the peripheral blood smear findings through manual microscopic examination. However, routine peripheral blood smear analysis is both time-consuming and labor-intensive. Here, we propose a smartphone-based autofluorescence microscopy (Smart-AM) system for imaging label-free blood smears at sub-cellular resolution and performing hematological analysis. Smart-AM enables rapid, high-quality, and label-free visualization of morphological features of different blood cells (leukocytes, erythrocytes, and thrombocytes) and abnormal variations in blood cells. Moreover, assisted with deep learning algorithms, this technique can automatically detect and classify different leukocytes with high accuracy, and transform the autofluorescence images into virtual Giemsa-stained images maintaining significant cellular features. The proposed technique is portable, cost-effective, and user-friendly, making it significant for broad point-of-care applications.

## 1. Introduction

Hematological analysis plays a pivotal role in clinical tests associated with the analysis of the cellular component of blood. Through the evaluation of the blood cell enumeration and morphology, hematological analysis enables screening and monitoring of blood-related conditions and diseases, like inflammation, anemia, infection, hemophilia, blood-clotting disorders, and several types of cancers. Hematological analysis has been conducted in many clinical applications. For example, in chemotherapy, the chemotherapy drugs will inevitably damage the normal cells of the human body while killing cancer cells due to their weak selectivity. It will result in adverse drug reactions (ADR) [1]. Myelosuppression [2], one of the most common adverse reactions of chemotherapy, is mainly manifested by thrombocytopenia and the decrease in the number of leukocytes (especially neutrophils) and hemoglobin. Hematological analysis can reflect whether the body has myelosuppression and the degree of myelosuppression. Hence, hematological analysis has been considered one of the most efficient tests before and after chemotherapy. Although flow cytometry and microscopic evaluation of stained blood smears are widely used in clinical laboratories and hospitals for hematological analysis, there remain some drawbacks. The whole procedure requires multiple chemical reagents and expensive microscopic machines, laborious system calibration, and highly trained personnel for operation. The disadvantages limit the analysis to being done in a hospital or clinical setting. However, going back and forth to the hospital twice a week for hematological analysis is annoying, especially for chemotherapy patients in fragile health. Thus, there is a great demand to develop a rapid, reagent-free, and portable imaging technique for hematological analysis.

Alternative novel imaging modalities have come out to achieve label-free hematological analysis. For instance, Raman spectroscopy has investigated leukocytes and visualized cellular components to differentiate leukocytes according to the different biomolecular information in the spectra [3,4]. However, the drawback is that the spontaneous Raman signal is weak for detection. Deep-ultraviolet (Deep-UV) microscopy [5] is another powerful emerging tool for molecular imaging without labels, and its contrast comes from biomolecules’ optical absorption properties. Deep-UV microscopy can achieve higher spatial resolution due to its short wavelength light for illumination and quantitative information from various endogenous biomolecules. Other techniques, such as hyperspectral microscopy [6–8], quantitative phase microscopy [9–13], and fluorescence lifetime microscopy [14], have been promising tools for cellular and subcellular imaging in biomedical fields. Although they show the enormous potential of characterizing changes in blood cell components for hematological analysis, the fact that they require complex and expensive optical devices with sophisticated system alignment makes non-professionals incompetent for the operation. In contrast, recent smartphones have shown great potential as a promising platform for point-of-care monitoring and diagnosing due to their widespread distribution. A considerable amount of studies have attempted to develop microscopy imaging modalities (e.g., bright-field, fluorescence, phase-contrast, etc.) based on smartphones with simple add-on lenses, especially for various medical diagnoses and patients’ medical record tracking [15–22].

Here, we propose a quantitative imaging technique, termed smartphone-based autofluorescence microscopy (Smart-AM) system, for imaging label-free blood smears at sub-cellular resolution and performing hematological analysis. A light-weight external lens module is attached to the typical smartphone’s built-in camera, and only small ultraviolet light-emitting diodes (UV-LEDs) are needed for illumination, which provides high-quality autofluorescence images with a maximum field of view of 2.58 mm × 1.94 mm. Smart-AM images can exhibit morphological features of different blood cells (leukocytes, erythrocytes, and thrombocytes) without labels and show abnormal variations in blood cells efficiently. Compared with the traditional hematological imaging system, our Smart-AM system is much more robust and cost-effective. Furthermore, we combine the imaging system with a novel detection and segmentation algorithm (e.g., Detectron2 [23]) and deep learning transformation networks (e.g., Pix2pix [24]), coined as DeepSmart-AM, to decrease the requirement for skilled hematologists to make the manual diagnosis. The performance of Smart-AM is experimentally verified by imaging mouse blood and human blood samples. The results show that specific features in Smart-AM images can be used for leukocyte differential automatically with high accuracy (over 90%), showing a high consistency with the manual counting results. Also, the deep learning transformation networks can convert the autofluorescence images into virtual Giemsa-stained images which are almost the same as the real Giemsa-stained blood smear images. This can simplify the complicated sample preparation procedures and decrease diagnosis errors caused by bad staining quality. This proposed technique simplifies the clinical workflow of the blood test from days to less than ten minutes, and eliminates the need for expensive machines, chemical reagents, and human labor, as shown in Figure 1b. It is portable, cost-effective, and user-friendly, making it significant for broad point-of-care applications.

**Figure 1.**
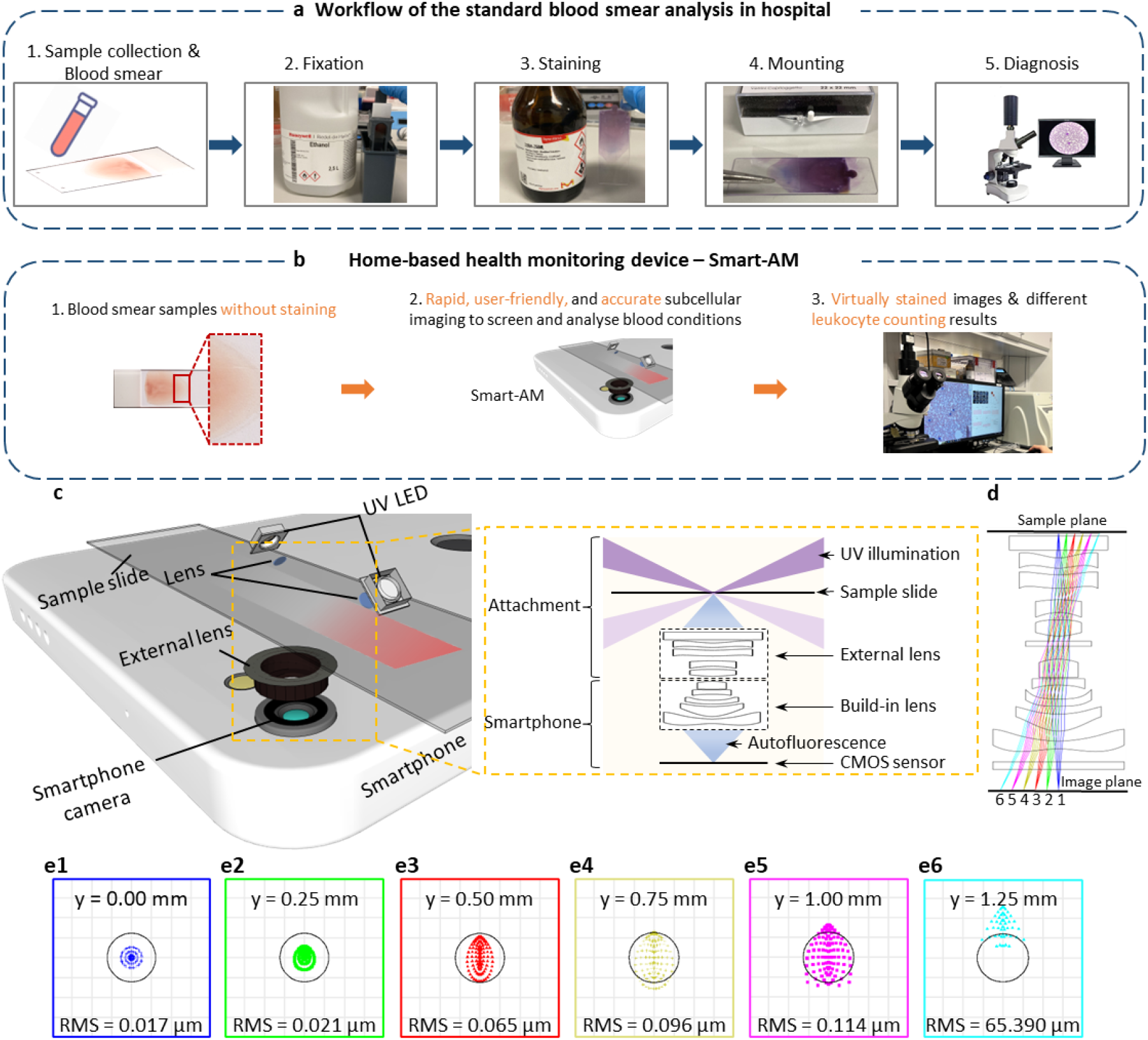
Motivation, design and simulation of Smart-AM. a) Conventional standard blood smear analysis. b) Proposed new home-based blood imaging and diagnosis by the Smart-AM system. c) Schematic illustration of Smart-AM system. The major components of Smart-AM system are ultraviolet light-emitting diodes (UV-LEDs), focus lenses, sample slide, an external lens module, and smartphone camera. The UV light is obliquely focused onto the sample by UV fused silica plano-convex lenses. The excited autofluorescence signal is collected by an external lens module, refocused by the build-in lens, and subsequently detected by the smartphone camera sensor. d) Ray-tracing simulation of the external reversed lens module served as an objective for smartphone-based microscope. e1–e6) Simulated spot diagrams at the image plane of six different fields (Δy = 0, 0.25, 0.50, 0.75, 1.00, and 1.25 mm from the center of the field at the sample plane). Minimal aberration was observed within the center 1 mm radius region.

## Materials and Methods

### Design of the Smartphone-based Autofluorescence Microscopy (Smart-AM) System

The setup of the Smart-AM system is illustrated in Figure 1c. Since previous studies suggested that many important endogenous fluorophores (e.g., hemoglobin, nicotinamide adenine dinucleotide hydride (NADH), cytoplasmic aromatic amino acids, etc.) can fluoresce with deep-UV excitation [25–27], small ultraviolet light-emitting diodes (UV LEDs, 265–270 nm center wavelength, 10–15 mW, 5.5–7.0 V, 100 mA, 3535 packages) are used to illuminate samples obliquely with a high incidence angle of ~70°. After welding the LEDs to the board, we test the actual wavelength profile (Figure S1, Supplementary Information) in the working voltage of LEDs using a spectrometer (Sarspec, Lda). The LEDs are focused on the sample plane by small UV Fused Silica Plano-convex lenses. The oblique illumination from the opposite direction can circumvent the use of fluorescence filters and reduce the background shadowing artifacts because the directly transmitted excitation light (light purple rays in the ray-tracing image, Figure 1c) are missed by the external lens. The excited autofluorescence signals are collected by a reversed camera lens module (harvested from another device’s camera replacement part) that serves as the objective lens. Then through the infinity-corrected internal lens in the smartphone’s rear camera, the object is finally imaged on the color complementary metal-oxide semiconductor (CMOS) sensor of the smartphone (iPhone 12 Pro, *f*/1.6, 1.4 μm photo sites, 12.2 mm^2^ sensor size).

Digital devices such as smartphones and laptops are usually equipped with camera lenses composed of multiple highly complex aspheric lenses, which minimize aberration and field curvature. Therefore, they are well-suited to optically match the smartphone as a miniature microscope. With the external lens, the whole smartphone-based system acts as a relay lens. The system forms the final image on the smartphone sensor plane when an object is positioned at the focal plane of the external lens. The optical magnification (*M*) can be calculated by *f*_1_/*f*_2_ theoretically, where *f_1_* and *f_2_* are the focal length of the phone’s internal camera lens and the external lens, respectively. However, accurate information about the outer dismantled lens is not easy to obtain, so we measure the magnification by imaging a specific line in the target (USAF 1951 Glass Slide Resolution Targets, Edmund) (Figure S2, Supplementary Information). The magnification ratio can be calculated as (*Num* × Δ*S_sensor_*)/*w*, where *Num* is the pixels of the line in the image, *ΔS_sensor_* is the pixel size of the smartphone camera sensor, and *w* is the real width of the line. According to the optical magnification, the effective pixel size 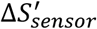 at the sample plane is Δ*S_sensor_*/*M*. Recent smartphone cameras have the advantage of a built-in lens with a high numerical aperture (NA) and an image sensor with a small pixel size, providing a high resolution for smartphone-based microscopy. According to the Rayleigh criteria, the theoretical resolution is approximately 0.61*λ*/*NA* = 2(*f*/#) × 0.61*λ*, where *λ* is the fluorescence emission wavelength, and *f/#* is the f-number of the external lens. Meanwhile, through Zemax optical ray-tracing and spot diagram simulation (six different fields, Δy = 0, 0.25, 0.50, 0.75, 1.00, *and* 1.25 *mm*, from the center of the field) (Figure 1e1–e6), minimal distortion, like coma, astigmatism, and field curvature, is observed within the center 1 mm radius region.

### Sample preparation

Mouse blood samples are collected by a terminal procedure (cardiac puncture), while human blood samples are collected from hospitals after the deidentification of patients. After collection, all blood samples are immediately added to vacuum tubes with the anticoagulant solution (Ethylenediaminetetraacetic acid solution, E8008, Sigma-Aldrich Inc.). Before making blood smears, the tube should be gently rolled to mix the blood cells. About 10 μL of whole blood is needed to drop on one end of the clean quartz slide. Another slide is used as the spreader to go back into the drop of blood at an angle of ~30–45 degrees. We push the spreader forward smoothly when we see the blood drop spreading along the edge of the spreader slide. A dense body, a well-developed feathered edge, and a monolayer area can be seen in a good blood smear. The monolayer of the blood smear is the main region we will use to execute the imaging tasks.

### Smart-AM and gold standard images data acquisition

The prepared blood smear is imaged directly using our Smart-AM system after air drying. As shown in Figure 1c, the blood smear is placed on the sample plane, which is between the reversed lens and illumination parts. A larger region of the blood smear can be imaged using a 2-axis translation stage for sample movement manually. The default smartphone camera application can be used for image acquisition directly, and other professional camera applications can also be used for advanced controls of exposure time, ISO, image format, etc. The focus alignment is easy due to the automated focus function of the smartphone camera. To impede the background light, a black box or cloth can be used to cover the imaging part.

After Smart-AM imaging, the same slide is manually stained to obtain the ground truth hematological image. Giemsa stain (32884, Sigma-Aldrich Inc.) is used, following the standard blood smear staining workflow (Figure 1a). The blood smear is first fixed in the methanol for 7 minutes and stained in a 1:20 diluted Giemsa solution for 30–60 minutes. After rinsing the stained smear with distilled water, the slide is allowed to dry in the air, and the permanent mount is made with the mounting medium. Then the slide is scanned by a digital whole-slide imaging machine (40×, NA = 0.75, NanoZoomer-SQ, Hamamatsu Photonics K.K.) to obtain corresponding Giemsa-stained hematological images. Note that the Giemsa-stained hematological images can duplicate the region imaged by the Smart-AM system with little distortion due to the fixation procedure.

### Automated, high-precision differential of five leukocytes with Detectron2 platform

Leukocytes have the most complex morphology among blood cells. Different leukocyte subtypes have different jobs, including recognizing intruders, killing harmful bacteria, and making antibodies. So, leukocyte identification is widely used to screen blood conditions and many diseases. Firstly, we used a manifold learning approach, locally linear embedding (LLE) [28], to visualize the autofluorescence image features (476 images). LLE is widely used in image recognition, high-dimensional data visualization, and other fields because it keeps the local features of samples in dimensionality reduction. In the next step, we can detect and differentiate leukocyte subtypes with relatively high accuracy by extracting the distinguishable characteristics of each subtype using an open-source detection platform with deep learning architecture (Detectron2 [23]). Detectron2 is implemented in PyTorch and includes numerous variants of the mask region-based convolution neural network (Mask R-CNN) model [29], which meet the segmentation and detection needs of leukocytes. The network combines feature map extraction, region proposals, bounding box regression, and classification of each RoI region. The Smart-AM images (image size of 256 × 256) of five leukocyte subtypes with corresponding type labels were used to train the network. In the testing phase, the input are Smart-AM images, and the output contains the bounding boxes, predicted leukocyte labels, and the confidence coefficient (the confidence thresh is setting to 0.75) of the predictions.

### Virtual staining of Smart-AM images with a conditional adversarial network

To show the effectiveness of Smart-AM in hematological imaging and to circumvent conventional chemical staining procedures, we have employed a conditional generative adversarial network (cGAN) from Pix2pix [24] in transforming our Smart-AM images to clinical standard Giemsa-stained hematology-like images (termed DeepSmart-AM images). Figure 6a illustrates the virtual staining training process using the Pix2pix algorithm, where paired Smart-AM images and Giemsa-stained blood smear images were used. The Smart-AM image is digitally transformed to the virtually Giemsa-stained image by the generator G. Then the discriminator D is used to classify between the generated virtually stained image and the actual stained image. The UNet-based G and PatchGan-based D are used in this work. We made two improvements on the original Pix2pix to force it to keep the structural information of Smart-AM images as much as possible. First, we add structure similarity index (SSIM) [30] loss that minimizes the structural differences between input Smart-AM images and generated Giemsa-stained images. In addition, we also train a generator F to transfer the Giemsa-stained blood smear images back to virtual Smart-AM images. With the F, the generated virtually stained image can be transferred back to Recovered Smart-AM images which can be used to calculate the recovering loss with original Smart-AM images. The recovering loss also forces the G to keep the structural information of Smart-AM images as much as possible. 1230 paired Smart-AM and Giemsa-stained image patches with a patch size of 256 × 256 (cropped from the 4036 × 3024 pixels original Smart-AM images with a 100-pixel moving step) were used in the training phase. The loss function of the network is defined as:

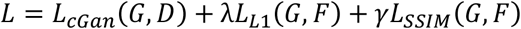

Here:

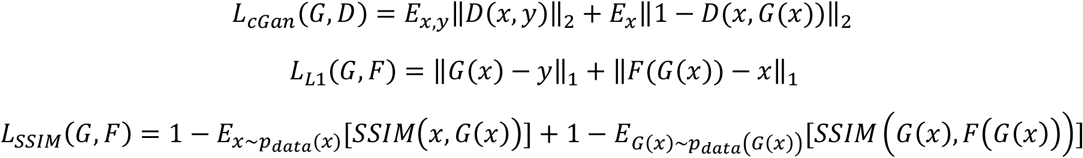

Where the *x* and *y* represent the input Smart-AM images and real Giemsa-stained images. We set the weighting parameters of the L1 loss λ = 10, and SSIM loss *γ* = 1. The optimal network is obtained through continuous optimization of the confrontation between the generator and discriminator. To demonstrate the possibilities and effectiveness of virtual staining Smart-AM (DeepSmart-AM) images, we imaged unstained blood smears using our Smart-AM system and then obtained the virtual Giemsa-stained images by the trained generator directly.

## Results

### Imaging performance of Smart-AM verified by fluorescent beads

The imaging resolution performance of our Smart-AM system was quantified by imaging green fluorescent polymer microspheres with a diameter of 200 nm (G200, Thermo Fisher) (Figure 2a). The data points of ten fluorescent beads were selected and averaged to measure the resolution by Gaussian fitting (Figure 2b). The full width at half maximum (FWHM) of the Gaussian fitting profile is 1.42 μm, representing the resolution of the Smart-AM system.

**Figure 2.**
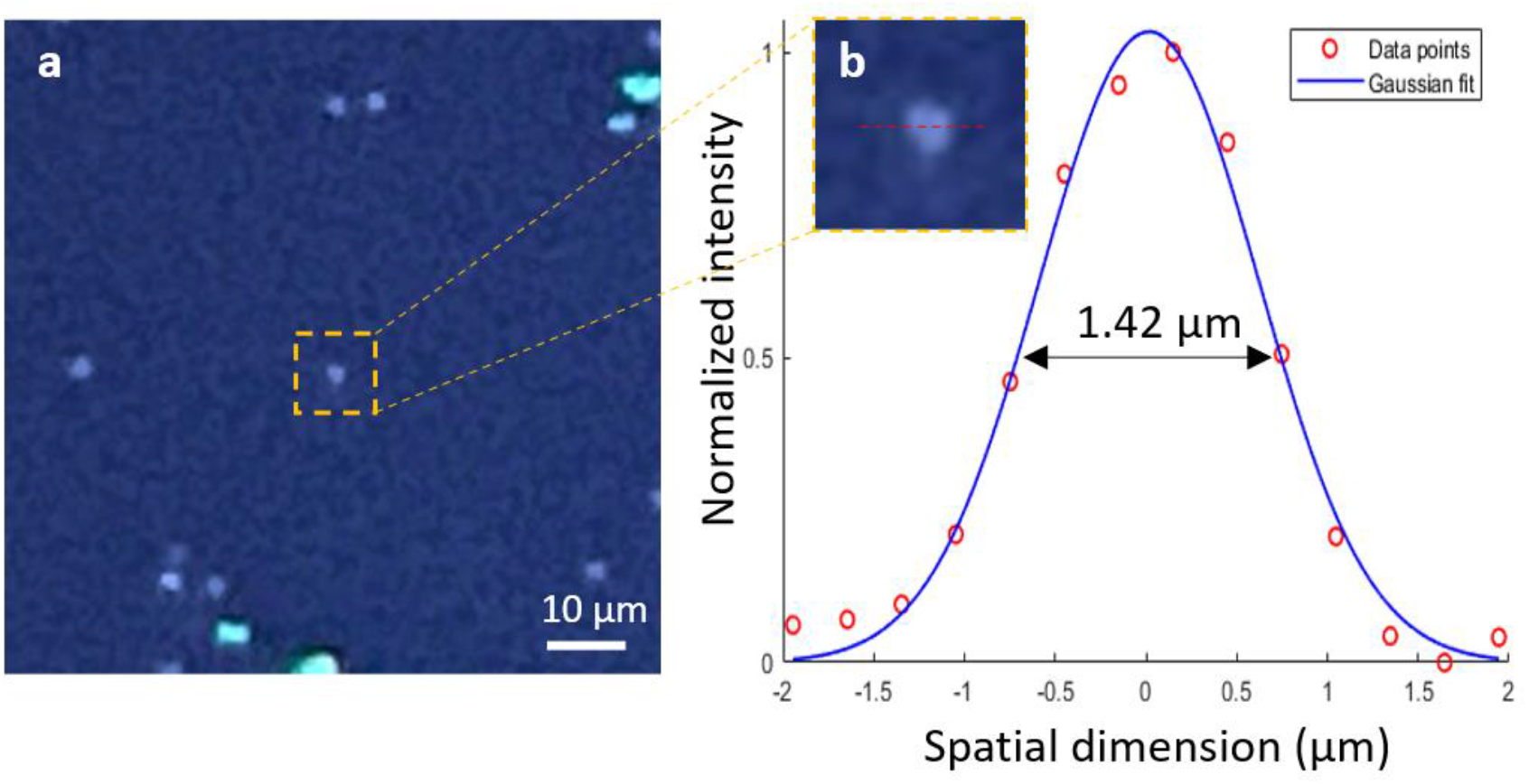
Smart-AM system resolution characterization. a) A Smart-AM image of fluorescent polymer microspheres of 200 nm in diameter. b) The zoomed-in view of one bead in the orange box in (a) and the profile along the red dashed line is extracted for averaging (red circles). The FWHM of the Gaussian fitting profile (solid line) is about 1.42 μm.

### Subcellular imaging of blood smears provided by Smart-AM

As shown in Figure 3, our system enables us to produce autofluorescence images of different blood cells. The first and the third columns of Figure 3 are Smart-AM images of the blood samples, and the second and the last columns are the corresponding Giemsa-stained images scanned by the digital whole-slide scanner. Again, the essential endogenous fluorophores can fluoresce under UV excitation, providing the autofluorescence imaging contrast of each cell type. Therefore, the recapitulate features of Smart-AM images show a significant consistency with the Giemsa-stained gold standard, which enables us to differentiate different blood cells, especially five different leukocytes. The UV absorption of nucleic acids in leukocytes produces distinguishable contrast in Smart-AM images, which corresponds to the violet color in Giemsa-stained images. The Smart-AM images for neutrophils (Figure 3a–c) present lobe textures of the nuclei (e.g., bilobed, trilobed, and multilobed neutrophils). Besides, eosinophils’ cytoplasm (Figure 3d,e, the white arrowheads) has stronger fluorescence compared to other granulocytes due to the rich acidophilic cytoplasmic granules.

**Figure 3.**
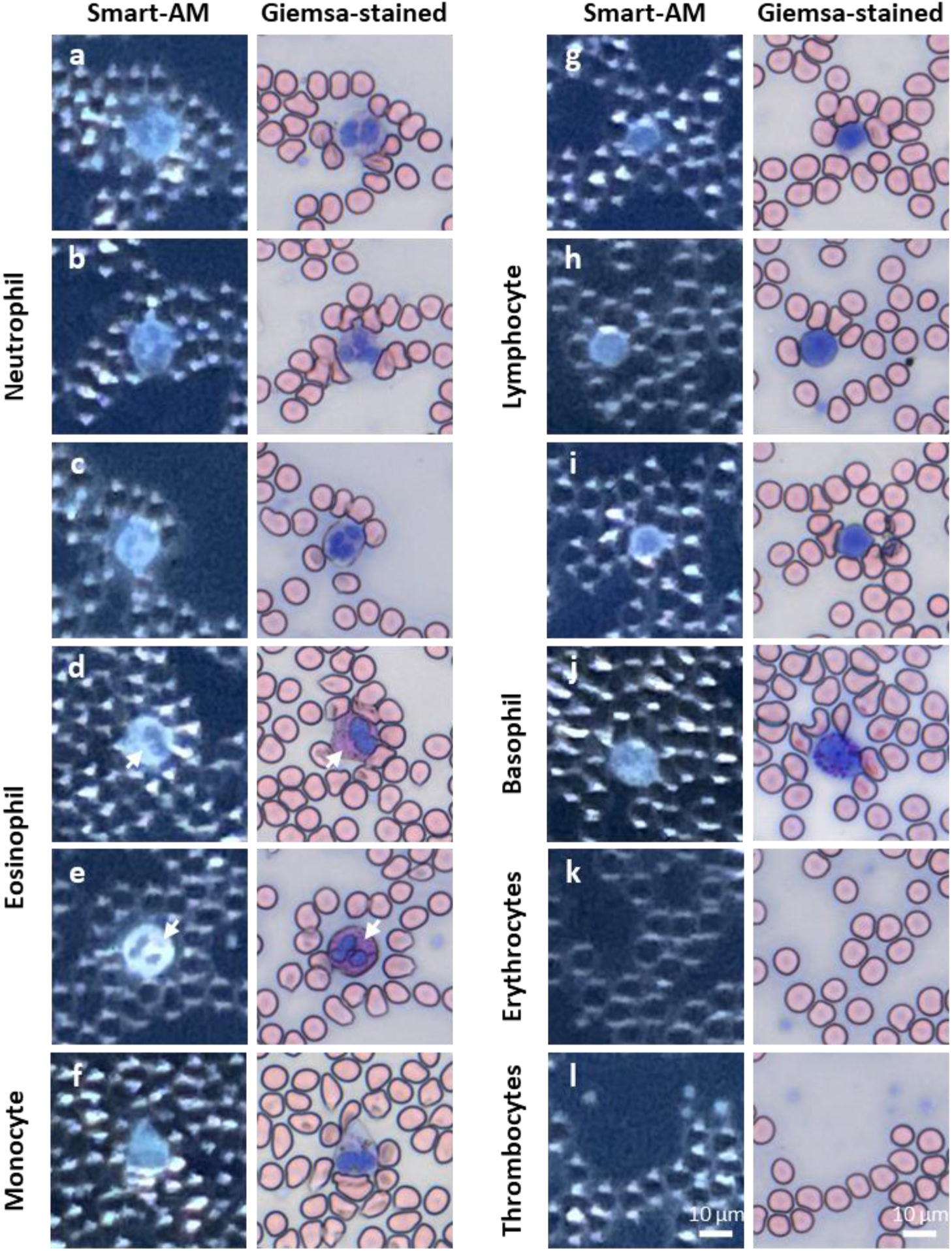
Smart-AM images and corresponding Giemsa-stained images of different blood cells. Different morphological details can be observed among different blood cells. a–c) Smart-AM and corresponding Giemsa-stained images of neutrophils. d,e) Smart-AM and corresponding Giemsa-stained images of eosinophil. f) Smart-AM and corresponding Giemsa-stained images of monocyte. g–i) Smart-AM and corresponding Giemsa-stained images of lymphocyte. j) Smart-AM and corresponding Giemsa-stained images of basophil. k) Smart-AM and corresponding Giemsa-stained images of erythrocytes. l) Smart-AM and corresponding Giemsa-stained images of thrombocytes.

### Atypical blood morphology visualized by Smart-AM

The Smart-AM images contain detailed characteristics which are essential for blood-related disease diagnosis and monitoring. Abnormal blood cell distribution or morphological structures can be observed in our Smart-AM images without sample processing or chemical reagents. To test this hypothesis, Smart-AM images from some abnormal blood samples were shown (Figure 4).

**Figure 4.**
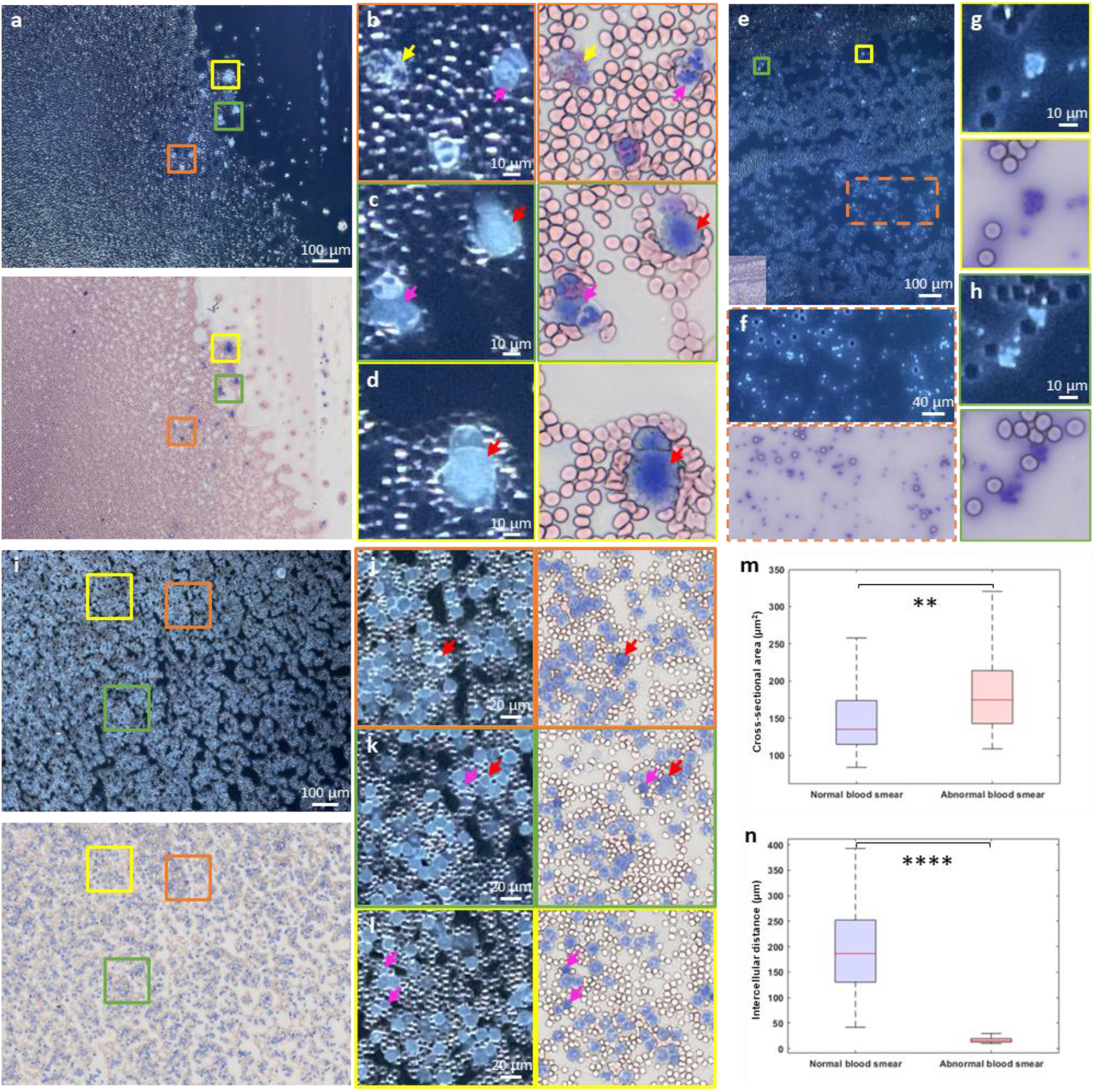
Smart-AM imaging of abnormal blood samples. a) Smart-AM (top) and corresponding Giemsa-stained (bottom) images over a ~1 mm × 1.4 mm region of a blood smear with abnormal leukocytes. b–d) Zoomed-in Smart-AM and Giemsa-stained images of orange, green, and yellow boxes marked in (a), respectively. There are some leukocytes under apoptosis, such as cytoplasmic blebbing on the surface (red arrows in (c), (d)), and apoptotic bodies from one eosinophil (yellow arrows in (c)). Besides, some clumps of leukocytes are observed (pink arrows in (d), (e)) due to inflammation or bacterial infections. e) Smart-AM images over a ~1 mm × 1.4 mm region of a blood smear with abnormal thrombocytes. The insert at the bottom left of the Smart-AM image is the corresponding Giemsa-stained image. f) Zoomed-in Smart-AM and corresponding Giemsa-stained image of orange dash region marked in (e), showing the excess platelets. g,h) Zoomed-in Smart-AM and corresponding Giemsa-stained images of yellow and green boxes marked in (e), respectively, showing platelets clumping. i) Smart-AM (top) and corresponding Giemsa-stained (bottom) images over a ~1 mm × 1.4 mm region of a blood smear with cancer. j–l) Zoomed-in Smart-AM and Giemsa-stained images of orange, green, and yellow boxes marked in (i), respectively. Excess neutrophils and larger lymphocytes (red arrows in (j), (k)) compared with normal lymphocytes (pink arrows in (k), (l)) are observed in Zoomed-in parts. m) Distribution of cross-sectional areas of leukocytes extracted from normal and cancer blood samples with a median of 134 μm^2^ and 175 μm^2^, respectively. n) Distribution of intercellular distances of leukocytes extracted from normal and cancer blood samples with a median of 186 μm and 17 μm, respectively.

Apoptosis [31] is usually a normal process of programmed cell death. However, apoptosis in both excessive and reduced amounts has pathological implications. Figure 4a–d shows excess leukocytes are under apoptosis (the red and yellow arrowheads). The red arrowheads indicate cytoplasmic blebbing on the surface, and the yellow arrowhead indicates one eosinophil is breaking into apoptotic bodies. Besides, some clumps of leukocytes (the pink arrowheads in Figure 4b,c) are observed due to inflammation or bacterial infections.

In the Smart-AM images of blood samples with platelet abnormalities (Figure 4e), large and excess platelets (zoomed-in Smart-AM image Figure 4f of the orange dash region marked in Figure 4e), and aggregate platelets (Figure 4g,h with corresponding Giemsa-stained hematological images) can be easily observed. These platelet abnormalities imply essential thrombocythemia or reactive thrombocytosis.

Figure 4i exhibits excess and abnormal leukocytes over a ~1.4 mm^2^ region. The zoomed-in Smart-AM images (Figure 4j–l) show the details of these leukocytes. The number of neutrophils increases dramatically in this sample, and the size of some lymphocytes (the red arrowheads) is bigger than normal lymphocytes (the pink arrowheads). Diagnostic features, such as cross-sectional area and intercellular distance of leukocytes, play a vital role in addressing the issue of blood conditions. These features are extracted from Smart-AM images of both cancer blood samples and normal blood samples by using a pixel-based segmentation Fiji plugin (trainable Weka segmentation [32]). The segmentation results can be analyzed directly in Fiji to get the area and position of each leukocyte. The statistical results (Figure 4m,n) indicate a significant difference between normal and cancer blood samples based on the cellular features.

Compared with Giemsa-stained images, the Smart-AM images can successfully identify atypical blood morphology and size, which verify the potential of our system for blood screening applications.

### Leukocyte five-part differential

Differential leukocyte counting is essential for hematological analysis to monitor and diagnose blood diseases. Here, we employed LLE to visualize the blood cell features in two-dimension and three-dimension spaces (Figure 5a). The result shows good clusterings between the blood cell types. To complete leukocyte subtype differential analysis, we employed an open-source platform, Detectron2, based on a deep learning algorithm. Some Smart-AM images of detection and classification results are shown in Figure 5c. The results contain the bounding boxes, predicted leukocyte labels, and the confidence coefficient. The confusion matrix represents the differential result by the counts of the predicted and actual class labels (Figure 5b). According to the matrix, the result was evaluated by using two quantitative metrics of accuracy and F1-score. The accuracy is the ratio of all correct predictions of instances to the whole pool of instances, which is the most intuitive metric. In contrast, F1-score is the harmonic average of the precision (the ratio of the correctly predicted positive labels to all the correct predictions) and recall (the ratio of the correctly predicted positive labels to the actual positive instances), which is better for our uneven class distribution. The average accuracy and F1-score among all leukocytes are 0.960 and 0.887, respectively. The differential performance of the monocyte and the basophil are weaker than other subtypes, showing that some monocytes and basophils are classified into neutrophils or lymphocytes due to some similar morphological features. This issue can be addressed by obtaining more evenly distributed samples or improving the deep learning algorithm. The Smart-AM images combined with the detection and classification deep learning network can provide the tool for leukocyte subtype differential with a favorable performance for hematological analysis.

**Figure 5.**
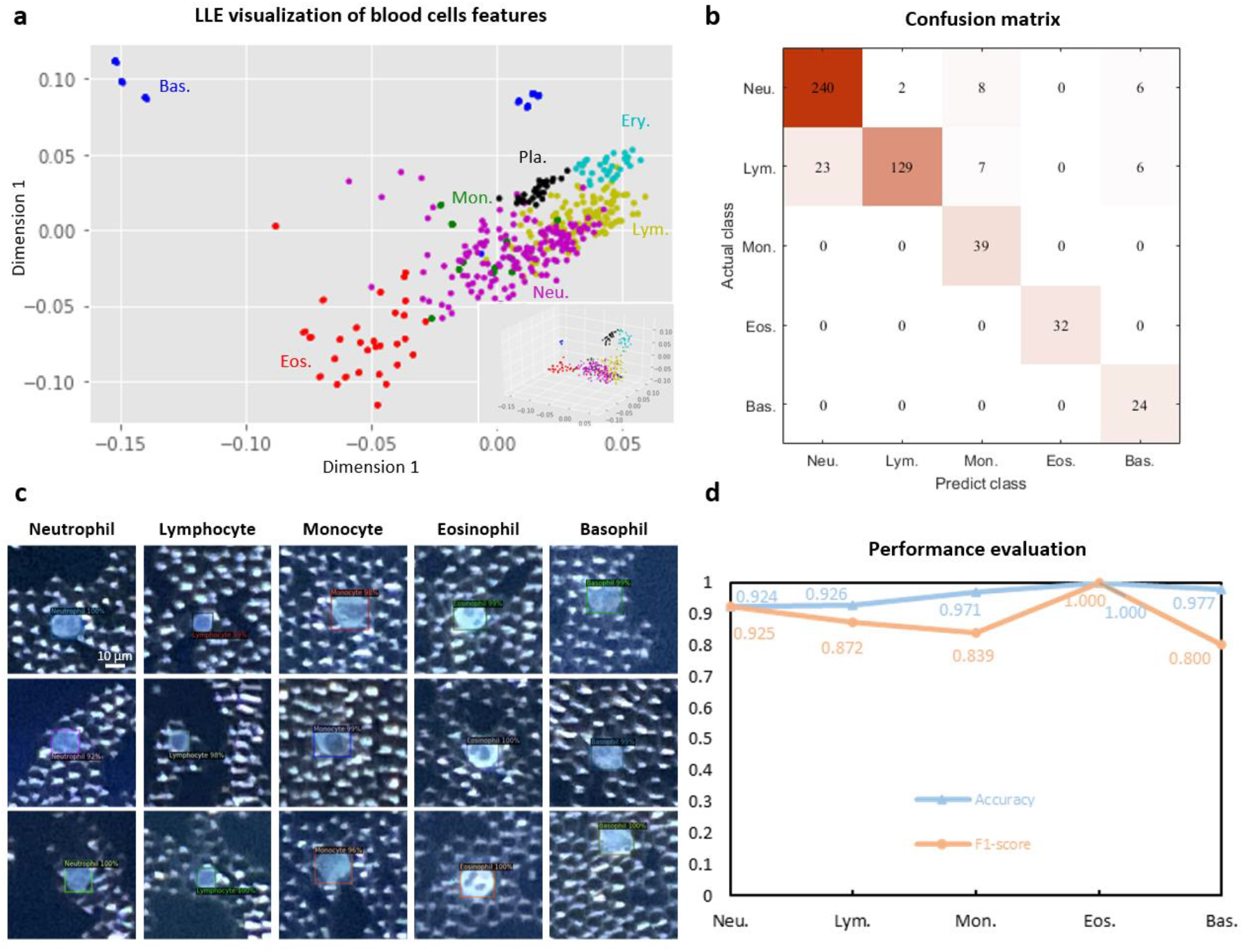
Automated, high-accuracy differential of five leukocytes with Detectron2 platform. a) The LLE visualization of blood cell features in two-dimension and three-dimension (inset at the bottom right) spaces. b) Confusion matrix for five-part leukocyte differential counts. c) Smart-AM images of five leukocyte subtypes from the detection results. The results contain the bounding boxes, predicted leukocyte labels, and the confidence coefficient. d) Performance evaluation of differential results by quantitative metrics (Accuracy and F1-score). The average values among all leukocytes are 0.960 and 0.887 for Accuracy and F1-score, respectively. Neu.: neutrophil, Lym.: lymphocyte, Mon.: monocyte, Eos.: eosinophil, Bas.: basophil.

To validate the performance of our Smart-AM images combined with the detection and classification deep learning network for screening leukocyte count disorders, we compared the leukocyte percentages from both our detection results and the manual counting results (Figure S3). Only slight differences are shown between predicted results and the manual counting results, which can be avoided using sufficient amounts of leukocytes. Meanwhile, from the results, our technique can provide a promising tool to show the abnormal variation in leukocyte percentages for screening patients’ blood conditions.

### DeepSmart-AM images of unstained blood smears versus traditional Giemsa-stained images

To make our Smart-AM a clinical translational technique, we transform the Smart-AM images into DeepSmart-AM images which mimic the appearance of real Giemsa-stained images and are more understandable for hematologists. Detailed information on the virtual staining network (Figure 6a) is shown in the Materials and Methods section. To demonstrate the clinical possibilities and effectiveness of our DeepSmart-AM images, we imaged fresh blood smears using our Smart-AM system and generated the DeepSmart-AM images by virtual staining network, then compared them with Giemsa-stained images of the same slides (Figure 6 and Video S1, S2, Supporting Information). Figure 6f–m shows Smart-AM images and their virtual staining version output (DeepSmart-AM images) of the zoomed-in regions in Figure 6b–e, and Figure 6n–u are the corresponding bright-field Giemsa-stained images as ground truth. Validated with erythrocytes, different leukocyte subtypes (Figure 6f–k), and platelets (Figure 6l,m), the well-trained network enables the transformation of Smart-AM contrast images of label-free blood smears into DeepSmart-AM images which contain conducive information for hematologists to analyze and diagnose. For example, the multilobes of the leukocyte nucleus can be observed in DeepSmart-AM images. The pink part of the eosinophil, which indicates the rich acidophilic cytoplasmic granules, is consistent with the clinical standard. To quantify the virtual staining results, we first segmented both DeepSmart-AM and clinical standard images to identify and localize each blood cell. Blood cells’ cross-sectional area and intercellular distance were extracted from both DeepSmart-AM and clinical standard images. We plotted the distribution of these features to find out the relationship between the DeepSmart-AM and clinical standard Giemsa-stained images (Figure 7). According to the Wilcoxon rank-sum test, the high *p* value indicates that it is hard to tell the differences between DeepSmart-AM and clinical standard images, meaning the style transformation is highly effective.

**Figure 6.**
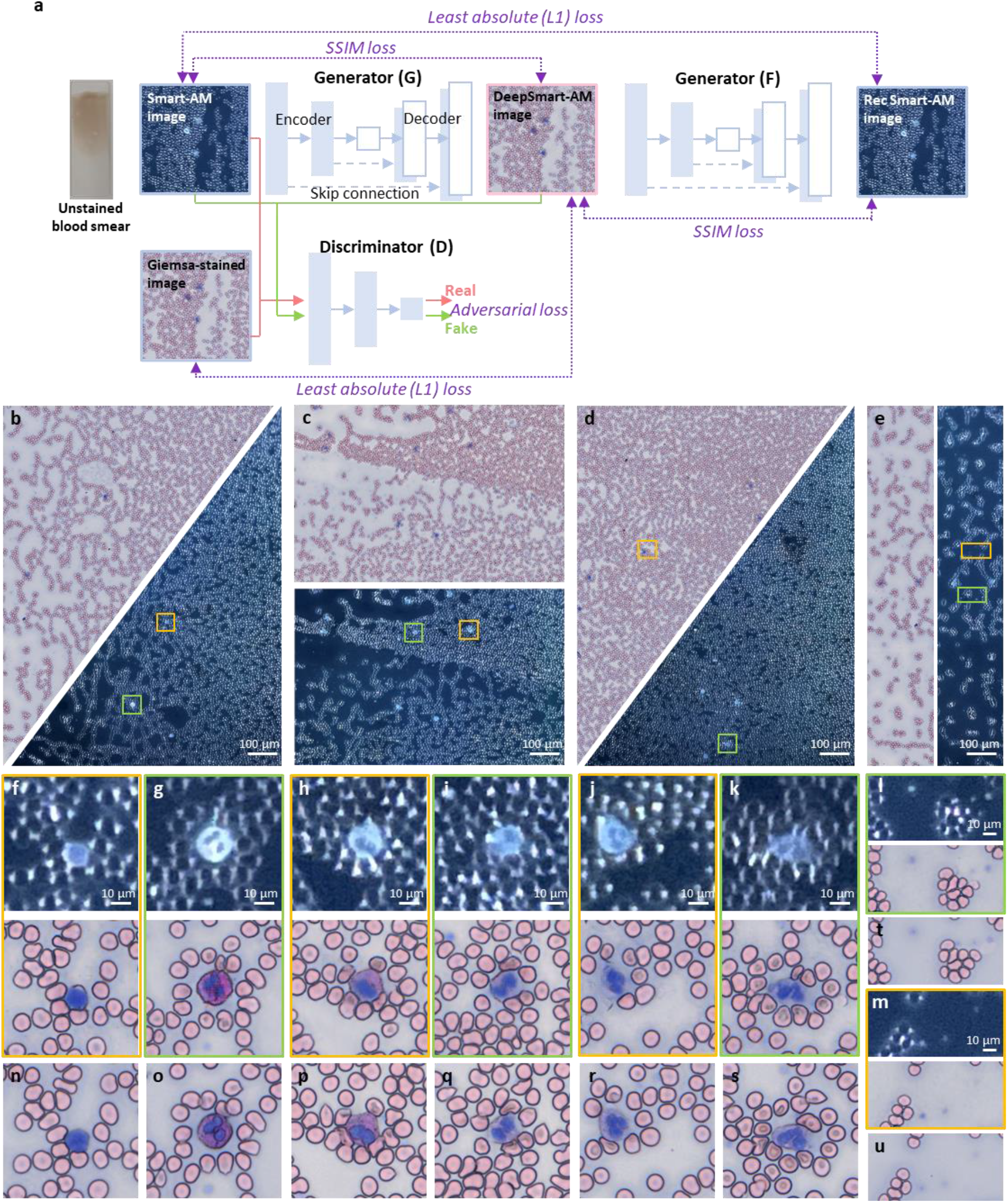
Virtual staining of the Smart-AM (DeepSmart-AM) images with a conditional adversarial network. a) Virtual staining by a conditional adversarial network with two generator and a discriminator. SSIM: structural similarity index measure, Rec: recovered. b–e) Smart-AM (bottom/right) and DeepSmart-AM (top/left) validation with blood smear samples. f–m) Zoomed-in Smart-AM and DeepSmart-AM images of orange and green solid boxes in (b–e), respectively. n–u) The corresponding Giemsa-stained images as the ground truth. Blood cells including erythrocytes, different leukocytes (lymphocyte in (f), eosinophils in (g) and (h), and neutrophils in (i–k)), and platelets, were well virtually stained to mimic the appearance of real Giemsa-stained images.

**Figure 7.**
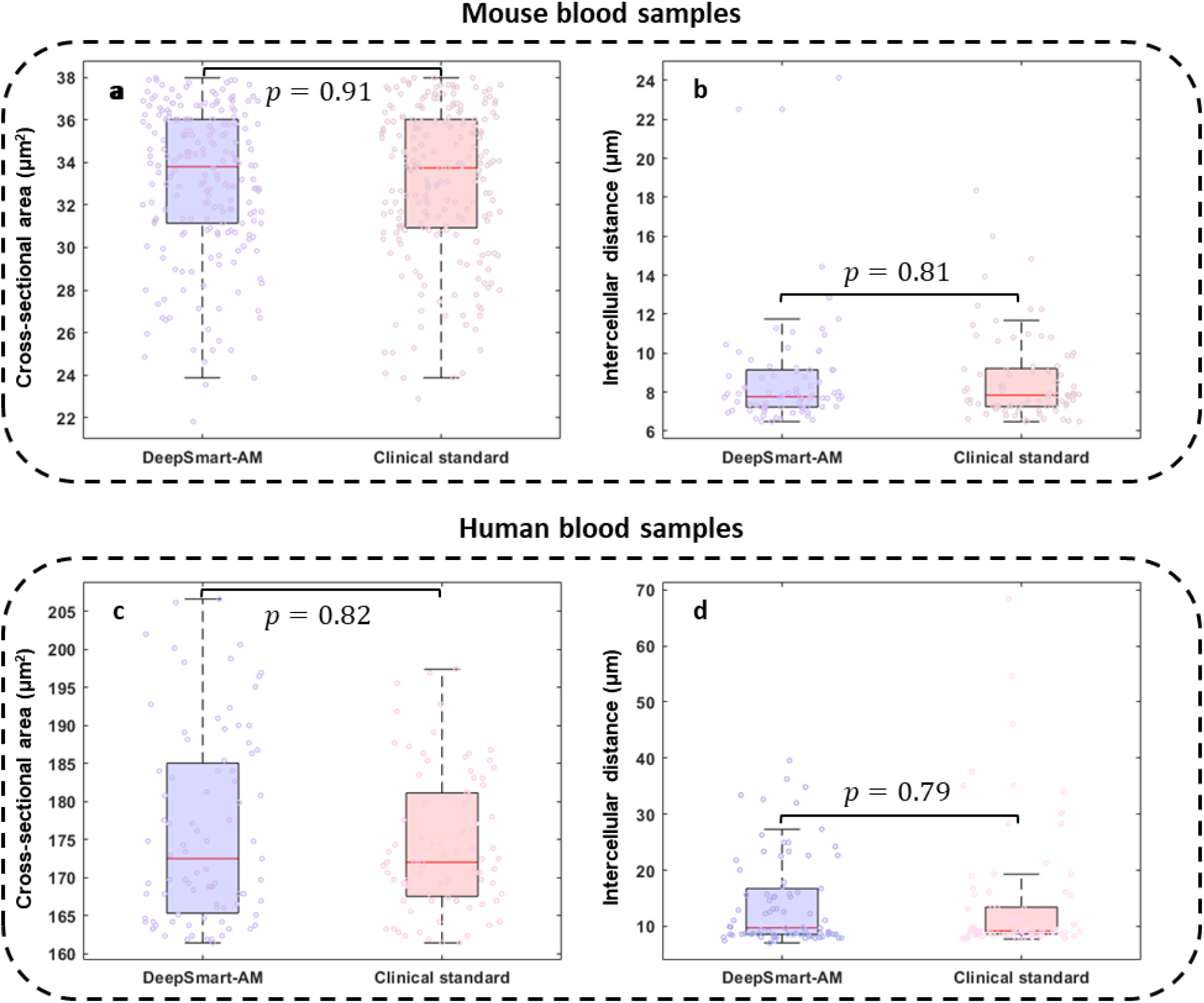
Distribution of blood cell features extracted from both DeepSmart-AM images and clinical standard Giemsa-stained images. a,c) Cross-sectional areas extracted from mouse blood samples and human blood samples, respectively. b,d) Intercellular distances extracted from mouse blood samples and human blood samples, respectively. Wilcoxon rank-sum testing is calculated for each distribution. The significance is defined as *p* ≤ 0.05, and the higher *p* value here means it is harder to tell the differences between the two images.

## Discussion

Our Smart-AM system, combined with deep learning algorithms, holds great promise as a home-monitoring device for blood smear imaging and diagnosis, providing great convenience to patients who suffer from blood-related diseases that require frequent blood tests. To the best of our knowledge, this work demonstrates the first smartphone-based autofluorescence microscopy for label-free hematological analysis. As fundamental evidence, blood morphology plays an integral role in the screening and diagnosing of blood abnormalities. In the traditional blood smear analysis workflow, the blood smear slide should be fixed, stained with chemical agents such as Giemsa, Wright-Giemsa stains, and carefully observed by professionals using bulky and expensive microscopy systems. Our technique can provide equivalent diagnostic information to the canonical method. The morphological features of blood cells can be visualized without exogenous agents due to the intrinsic fluorescence contrast using UV excitation.

Meanwhile, the cost of our system (approximately no more than $100 for add-on parts and $1000 for the smartphone) is markedly cheaper than the commercial machines combined with multiple agents for hematological analysis. The deep learning-based detection and transformation algorithms enable rapid, practical, and reliable visualization and characterization of peripheral blood smears for non-professional users. These attributes make it translational to clinical settings and significant for broad point-of-care applications.

Despite successful demonstrations, further improvements still need to be conducted to meet some challenges and make our technique more applicable for clinical translations. First, the current system is a transmission mode for blood smears imaging, which limits its applications to thin slides and monolayer imaging. A reflection mode Smart-AM system can be developed through the total internal reflection field (TIRF) illumination method [33] or by using optical fibers to guide the illumination beam from the same side as the detection beam. In this way, the system can be applied to image larger regions of the blood smears that are not monolayer, or to image blood samples in blood bags directly without smearing. Second, a smartphone with a large sensor and small pixel size, combined with an external lens module with a shorter focal length, can be used to improve effective resolution and magnification. Third, some unsupervised deep learning methods can also be explored to impede the registration procedure in the network training part and enhance the detection and transformation efficiency. Finally, our DeepSmart-AM approach has focused on the normal mouse and human blood samples in this manuscript. More investigations are expected on the detection and style transformation of various abnormal blood samples.

In summary, we developed a smartphone-based quantitative autofluorescence microscopy system assisted with the deep-learning algorithms to achieve label-free blood smear imaging for hematological analysis, including visualization of essential features in blood cells, accurate detection and classification of different leukocytes, and high-quality virtual Giemsa-stained image transformation. This practical technique simplifies the traditional clinical workflow of the blood test from days to less than ten minutes and circumvents expensive machines, chemical agents, and human labor. It has great potential to enable point-of-care monitoring and remote hematological diagnostics in the future.

## Data availability

Data underlying the results presented in this paper are not publicly available at this time but may be obtained from the authors upon reasonable request.

## Supplemental document

See Supplement 1 for supporting content.

